# Contextualising transcription factor binding during embryogenesis using natural sequence variation

**DOI:** 10.1101/2024.10.24.619975

**Authors:** Olga M. Sigalova, Mattia Forneris, Frosina Stojanovska, Bingqing Zhao, Rebecca R. Viales, Adam Rabinowitz, Fayrouz Hamal, Benoît Ballester, Judith B Zaugg, Eileen E.M. Furlong

## Abstract

Understanding how genetic variation impacts transcription factor (TF) binding remains a major challenge, limiting our ability to model disease-associated variants. Here, we used a highly controlled system of F1 crosses with extensive genetic diversity to profile allele-specific binding of four TFs at several embryonic time-points, using *Drosophila* as a model. Using a combined haplotype test, we identified 9-18% of TF bound regions impacted by genetic variation. By expanding WASP (a tool for allele-specific read mapping) to examine INDELs, we increased detection of allele imbalanced (AI) peaks by 30-50%. This fine-grained ‘mutagenesis’ could reconstruct functionalized binding motifs of all factors. To prioritise potential causal variants, we trained a convolutional neural network (Basenji) to predict TF binding from DNA sequence. The model could accurately predict experimental AI for strong effect variants, providing a mechanistic interpretation for how genetic variation impacted TF binding. This revealed unexpected relationships between TFs, including potential cooperative pairs, and mechanisms of tissue specific recruitment of the ubiquitous factor CTCF.

## INTRODUCTION

The majority of disease-associated genetic variants occur in the non-coding genome (Edwards et al. 2013; Buniello et al. 2019), and likely affect gene regulation (Claringbould and Zaugg 2021). One mechanism by which mutations impact transcriptional programs is through the disruptions of transcription factor (TF) binding sites within regulatory elements, such as enhancers and promoters (Wray 2007; Fairfax et al. 2012; Abramov et al. 2021; Chen et al. 2016; Floc’hlay et al. 2021; Cannavò et al. 2016). However, establishing a mechanistic link between DNA sequence variation and TF binding remains challenging (Deplancke et al. 2016). Sequence variations that disrupt a TF’s cognate motif can only explain a minority of variable TF binding (Ding et al. 2014; Reddy et al. 2012; Kasowski et al. 2010). For example, for 24 TFs, only 12% of variable TF occupancy can be explained by variants in the TF’s motifs in a human lymphoblastoid cell line (Reddy et al. 2012). This implies that more complex mechanisms are involved in the majority of cases, including the presence of cryptic low affinity binding sites (Kribelbauer et al. 2019; Crocker et al. 2015), cooperative binding (Ibarra et al. 2020; Jolma et al. 2015; Spitz and Furlong 2012), and long-range interactions (Rao et al. 2014). In cases where TFs bind to DNA cooperatively (Jolma et al, 2015; Spitz & Furlong, 2012b), disruption of one of the two TFs’ motif can lead to a loss of binding of the second factor, even though its motif remains intact. At least 7.5% of allele-specific changes in PU.1 binding, for example, could be explained by genetic variation in four additional motifs (NFKB1, POU2F2, PRDM1, and STAT2) located in proximity to the PU.1-bound sites (Kilpinen et al, 2013; Raza et al, 2014; Heinz et al, 2013). For some cooperative TF::TF pairs, the binding site is different or a composite of the motifs for the individual TFs (Jolma et al. 2015), or may be driven by DNA shape rather than sequence (Ibarra et al. 2020), and therefore difficult to predict a priori. For other factors, variants that completely disrupt the TF’s motif can have no impact on their occupancy, as they remain recruited to the enhancer through protein-protein interactions as a TF collective (Khoueiry et al. 2017; Junion et al. 2012). In addition, the ability of some TFs to bind to DNA is altered by changes in chromatin accessibility, e.g. through changes in nucleosome occupancy (Barozzi et al. 2014). Thus, cooperative TF::TF interactions, as well as local TF::chromatin interactions make it difficult to infer the effects of genetic variation based on sequence data alone.

An additional challenge is the confounding effects of both *trans* and *cis* variation (Hill et al. 2021; Van De Geijn et al. 2015). Differences in TF binding across a group of genetically diverse individuals can be due to the impact of genetic variation on either (or both) the target DNA regulatory sequence (*cis*) or to mutations in the TF’s protein sequence, altering its binding affinity or specificity (*trans* effect). Allele-specific analyses is a very powerful method to tease apart *cis* and *trans* effects, i.e. by considering the ratio between two alleles in heterozygous positions thus normalising for differences in *trans* factors across individuals. Allele-specific analysis can thereby determine the functional impact of genetic variants to molecular phenotypes in *cis*, and provides an excellent ‘mutagenesis’ system to link genotype to phenotype in a quantitative controlled manner. Information on allele-specific TF binding can thereby greatly improve functional annotation of non-coding genetic variants associated with disease (Maurano et al. 2012; Walters et al. 2023; Cavalli et al. 2016), as shown in human lymphoblastoid cell lines (Kilpinen et al. 2013; Tehranchi et al. 2016) and patient tissues (Cavalli et al. 2016). In addition to pinpointing direct *cis* effect within the TF’s motif, F1 studies can also help to contextualise how TF’s function with other factors within their endogenous regulatory element (enhancer, promoter), as shown for three liver-specific TFs in mouse F1 hybrids (Wong et al. 2017) and three erythroid TFs in human cell lines (Behera et al. 2018), revealing new potential cooperative TF pairs.

Here, we systematically assessed the effects of *cis* genetic variation (both SNPs and INDELs) on TF binding during embryonic development, using genetic variation as a perturbation tool to better understand the sequence requirements for the recruitment of four essential TFs during embryogenesis. This includes three lineage factors involved in the subdivision and specification of the myogenic mesoderm (Twist, Mef2, Biniou) and one ubiquitous factor involved in chromatin topology (CTCF). By generating eight F1 crosses, we analysed allele-specific binding of the four TFs at multiple embryonic time-points (6 conditions in total), leading to a total of 60 ChIP-seq datasets, each with biological replicates (120 experiments). To take advantage of INDELs in addition to SNPs, we expanded WASP (Van De Geijn et al. 2015) to account for potential mappability biases of small deletions, and applied a combined haplotype test to identify allelic imbalance (AI). This uncovered 313-843 TF peaks (9-18% of each condition’s peaks set) that are affected by genetic variation in the context of embryogenesis, which is substantial given that three of these factors are essential regulators of development. By collating information from many functional variants that impact TF binding sites *in vivo*, these AI preferences were sufficient to reconstruct motif binding affinity. To pinpoint the mechanism of ‘hidden’ variation in TF binding, we trained a deep neural model (Basenji) on ∼1,200 ChIP-seq (Hammal et al. 2022) and DNAse (Reddington et al. 2020) tracks from embryonic and adult *Drosophila melanogaster* samples. The trained Basenji model, which is provided as a resource for future studies, can accurately predict occupancy from sequence in the reference genome, and when applied to the F1 data could infer allelic-effects of variants on TF binding. Notably, the model could predict the direction of strong effect variants across all TFs with an accuracy >90%, and revealed the relevant cognate and co-factor motifs, elucidating the causal mechanisms for a subset of variants. In some cases, this uncovered cooperative or dependent relationships between TFs, many of which were unknown, while in other cases it pinpointed a single base between two motifs, suggesting that DNA shape may play a role.

Taken together, this study provides a comprehensive characterization of the sequences affecting TF binding in the context of embryonic development, and provides a quantitative and global assessment of functional bases required for the occupancy of these essential developmental regulators.

## RESULTS

### Quantifying TF binding across genetically diverse embryos

We generated an extensive dataset of chromatin immunoprecipitation followed by sequencing (ChIP-seq) for four TFs (Twist, CTCF, Mef2, and Biniou) at several time-points during embryonic development in eight F1 crosses of inbred lines of *Drosophila melanogaster* and two parental lines (Fig. 1A-B, Supplementary Table S1). All F1s crosses share the same maternal line (‘virginizer line’, referred to as VGN), providing a controlled genetic background. The eight paternal lines come from the *Drosophila* Genetic Reference Panel (Huang et al. 2014) and have a high level of genetic diversity among them. Three of the analysed TFs (Twist, Mef2, and Biniou) form a gene regulatory network that is essential for mesoderm specification and its derived muscle tissues’ development, and have a hierarchical relationship between each other (Zinzen et al. 2009; Jakobsen et al. 2007; Sandmann et al. 2007), allowing us to address the effects of combinatorial TF binding. The fourth factor, CTCF, is involved in 3D chromatin structure and potentially other roles in gene expression (Narendra et al. 2016; Gambetta and Furlong 2018; Li et al. 2022; Dehingia et al. 2022). All four TFs have different types of DNA binding domains: bHLH (Twist), MADS-box (Mef2), forkhead (Biniou) and C2H2 zinc finger (CTCF), and therefore recognise DNA in different ways, which may influence the impact of genetic variation. Moreover, all four factors are conserved from flies to humans, with highly conserved DNA binding domains, implying that the sequence rules learnt here will likely be conserved in other species.

**Figure 1.**
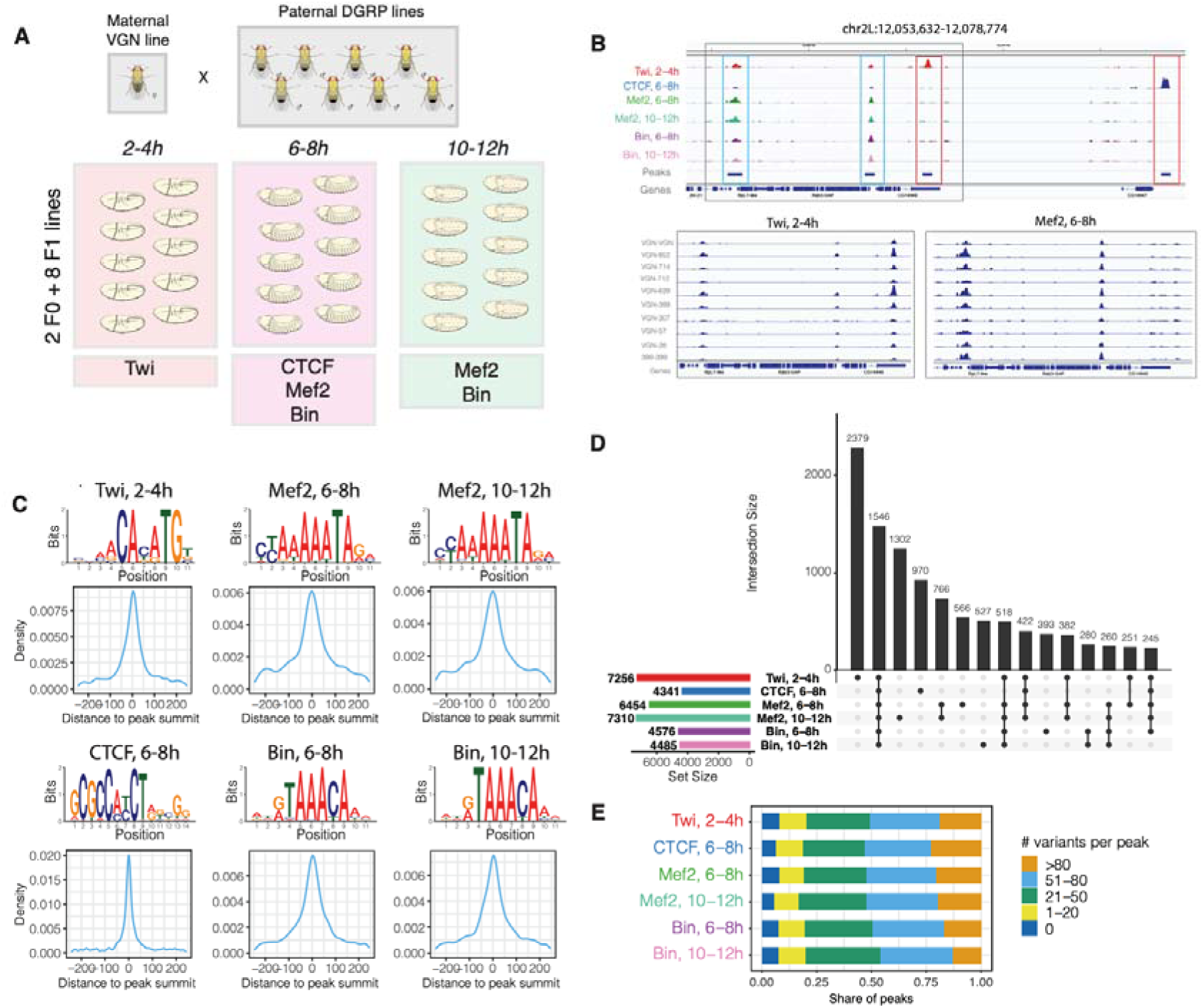
Profiling TF binding in F1 embryos of *Drosophila melanogaster*. **A.** Experimental design: F1 crosses were generated between maternal VGN line and eight paternal DGRP lines. Staged embryos were collected at three timepoints (2-4h, 6-8h, 10-12h). Binding of four TFs (Twi, CTCF, Mef2, and Bin) was profiled at the indicated embryonic time-points, in two parental lines (VGN and DGRP399) and eight F1 lines. **B.** *Top*: Occupancy of all TFs in the chromosomal region chr2L:12,053,632-12,078,774. For each TF, tracks were generated by averaging signal across all ten lines at the corresponding time-point. Blue boxes indicate regions co-bound by multiple factors, red boxes show peaks specific to one TF. *Bottom*: binding of Twi at 2-4h (left) and Mef2 at 6-8h (right) across the individual lines on chr2L:12,054,806:12,066,910 (region marked as grey box on the top plot). **C.** *De novo* motifs discovered in top-1000 ChIP-seq peaks (top) and central enrichment of these motifs in full consensus peak sets of Twi at 2-4h, CTCF at 6-8h, Mef2 at 6-8h, Mef2 at 10-12h, Bin at 6-8h, and Bin at 10-12h. **D**. Overlaps among consensus peak sets for all TFs.

As part of the DGRP project, the parental lines were obtained as wild isolates and inbred for over 20 generations. They are close to homozygosity (∼99.7%) and have extensive genetic variation between them (Huang et al. 2014). We genotyped all parental lines to tailor the variant call to our in-house versions of these fly lines (which necessarily went through further inbreeding and bottlenecks in the years following the original genotyping) to increase sensitivity, by providing 74-125X coverage of the *D. melanogaster* genome across each line (Methods). Variants were called applying the GATK4 (Poplin et al. 2018) best practice pipeline and the results filtered with two sets of thresholds leading to a stringent variant set used in the association tests (that minimises false discovery) and a lenient variant set used to correct for mappability issues (that minimises false negatives). A total of 1,735,077 unique heterozygous positions were identified in at least one of the F1 individuals (96% bi-allelic variants) in the stringent variant set (1,912,835 in the lenient), thereby allowing for a genome wide assessment of the impact of genetic variation on TF binding.

Tightly staged embryo collections, centred on the embryonic stages of key functional importance for each TF, were used for the ChIP-seq experiments (Fig. 1A). For Twist, this was during early embryogenesis (2-4h, predominantly stage 5-6), where *twist* is required for mesoderm gastrulation and its subsequent division. *Mef2* and *biniou* have broad expression, spanning many embryonic stages and are required for the specification and differentiation of somatic (Mef2) and visceral (*Mef2*, *biniou*) muscle. We therefore profiled both TFs at two stages: 6-8h (predominantly stage 10/11) during cell fate specification, and at 10-12h (predominantly stage 13) at the initiation of tissue differentiation (Fig. 1A). This allows for the detection of condition specific effects of genetic variants for the same factor – constitutive (both stages) or only one of the two stages. The vast majority of sites occupied by CTCF don’t change during embryogenesis, or even between tissues (Pollex et al. 2024), and we therefore profiled one representative timepoint at mid-embryogenesis (6-8h).

When mapping ChIP-seq reads from different genetic backgrounds, mapping biases need to be considered. The state-of-the-art tool for handling mapping bias, WASP (Van De Geijn et al. 2015), discards INDELs. However, since INDELs span multiple base-pairs (bp), they are intuitively more likely to have a stronger impact on TF binding compared to single nucleotide polymorphisms (SNPs) when overlapping a TF motif. In addition, masking INDELs can cause spurious associations in the case of bystander SNPs that are in linkage with causal INDELs. The majority of INDELs in these F1 embryos are short (70% below 5 bp, 87% below 10 bp) and as the ChIP-seq was performed with paired-end (PE) sequencing, we reasoned that including INDELs should improve the power to detect the impact of genetic variation without creating mappability issues. Relatively long read paired-end sequencing (81bp PE for ChIP samples, 151bp PE for ChIP inputs) also raises the proportion of reads overlapping INDELs, thus increasing the number of reads that would be discarded if INDELs were ignored (Supplementary Fig. 1A-B). To address these issues, we made significant modifications to the original WASP codebase to now enable reads to overlap INDELs. We included further additional options that speed up quantification of genetic variants over fixed intervals (e.g. TF peaks), and increased power to detect errors coming from incorrect genotyping (Methods, https://git.embl.de/rabinowi/wasp_indels). This new version of WASP (WASP-INDEL) applies the same algorithm to both SNPs and INDELs making it equally effective in removing mapping biases caused by either. This increased the usable reads by ∼13-28% compared to the original software and allowed us to test for the effect of 125,432 INDELs on TF binding present in our F1 crosses, leading to a total of 1,860,509 variants (SNPs and INDELs) that could be tested (+7,2% of tests).

After correcting for mapping bias, the reads were used for peak calling applying irreproducibility discovery rate (IDR) analysis (Methods). The majority of samples (96% (116/120)) showed high reproducibility between biological replicates (Methods, Supplementary Table S2). One replicate of line VGN-DGRP714 failed for both Bin and Mef2 at 10-12h, and the corresponding samples were excluded from further analyses. We constructed a consensus peak set for each condition (TF/time-point) using the DiffBind package (Stark R 2011) (Methods) by requiring peaks to be present (1bp overlap) in F1 embryos from at least three different lines (1% IDR, conservative set) and recalculating summits for each peak based on counts from all samples. This identified 7,256 peaks for Twist 2-4h, 6,454 for Mef2 at 6-8h and 7,310 at 10-12h, 4,576 for Biniou at 6-8h and 4,485 at 10-12h, and 4,341 for CTCF at 6-8h (Fig. 1D). Overall, all lines showed a high correlation between biological replicates over the consensus peak sets, attesting to the quality of the data (Supplementary Fig. 2A). Moreover, *de novo* motif discovery (Bailey et al. 2009) identified the expected motifs for all TFs, which showed strong central enrichment around the consensus peak summits (Fig. 1C, Methods). The majority of TF peaks in all conditions (87-96%, Supplementary Fig. 2B) were found in regions of open chromatin, defined by DNase-seq at the same stages of embryogenesis (Reddington et al. 2020).

The different TFs had highly overlapping binding patterns (considering peak summits within 500 bp), with 11% (1,546/13,693) of the combined consensus peak set present in all 6 conditions (Fig. 1D). While this could be expected for TFs involved in the same tissue’s development (Twist, Mef2, Biniou), the high overlap of CTCF at these putative enhancers was surprising. 55% of regions are co-occupied by at least two TFs (Fig. 1D). Sites co-bound by many TFs are enriched among TSS-proximal (<500bp from an annotated TSS) DHS and constitutively accessible distal DHS sites, while sites bound by single TFs are more distal and condition-specific, and likely putative condition-specific enhancers (Supplementary Fig. 2C-D).

Due to the extensive genetic diversity among these inbred wild-isolates, which have a median of 1 SNP per 227-236bp, around 88% of TF peaks have at least one genetic variant within 2.5 kB of the peak summit and 82% have more than 20 variants (Fig. 1E). As expected, TF binding for each factor was highly variable across genotypes (Fig. 1B, bottom).

### Effects of genetic variation on TF binding are extensive and preferably detected in genomic regions with less buffering

To disentangle the effects of genetic variation on TF binding from the other potential sources, we took advantage of the F1 experimental design to focus on allele-specific binding, which increases power to detect *cis* effects (McManus et al. 2010; Wong et al. 2017; Link et al. 2018; Floc’hlay et al. 2021). Specifically, we applied a combined haplotype test (CHT) (Van De Geijn et al. 2015), which simultaneously tests for the effect of each variant on read depth across different genotypes (beta negative-binomial (BNB) component) and for allele imbalance in heterozygous lines (allele-specific (AS) component) using a maximum likelihood approach (Methods). Total and allele-specific read counts were quantified per peak in each of the six consensus peak sets. Using crosses of highly inbred parental lines enables fully resolved haplotype phasing, which allowed us to test variants located ±2,5kb from the target TF peaks for association, resulting in a total of 475,1032 autosomal variants, including 45,597 INDELs). Importantly, the distribution of allelic ratios (share of reads mapped to one of the two alleles) was highly consistent across samples and centred around 0.5 (Supplementary Fig. 3A), which is the expected allelic ratio under null hypothesis in CHT (no allelic imbalance).

We identified strong genetic effects on the occupancy of all TFs, with the number of significant variants (FDR<1%; minimum allele imbalance AI=0.1) ranging between 4,264 for Biniou at 10-12h and 13,642 for Twist at 2-4h (Fig. 2C). Given the relatively low number of samples in our dataset, the allele-specific component of the test (CHT AS, Fig. 2C) had more power than the read depth component (CHT BNB, Fig. 2C) to detect genetic effects, resulting in stronger deviation of actual p-values from the uniform distribution and higher number of significant variants. Combining the two components (CHT full, Fig. 2C) resulted in slightly lower numbers of significant variants compared to the AS component alone, but provides a more stringent set of significant variants by adding constraints on the total read counts. Permuting genotype information also revealed some deviation from the uniform p-value distribution (CHT permuted, Fig. 2C). This likely results from the close relatedness among lines (all F1s sharing the same maternal line), which makes it impossible to shuffle genotype information for some variants (also discussed in (Van De Geijn et al. 2015)). However, given the stringent threshold on significant variants, almost all genetic associations were lost upon permuting genotype information, with only 12-255 significant variants remaining per condition. These numbers fall within the CHT FDR threshold, with a proportion of permuted genotype associations of 0.28-2.0%, indicating that the relatedness of the fly lines does not lead to an excess of false associations in this study.

**Figure 2.**
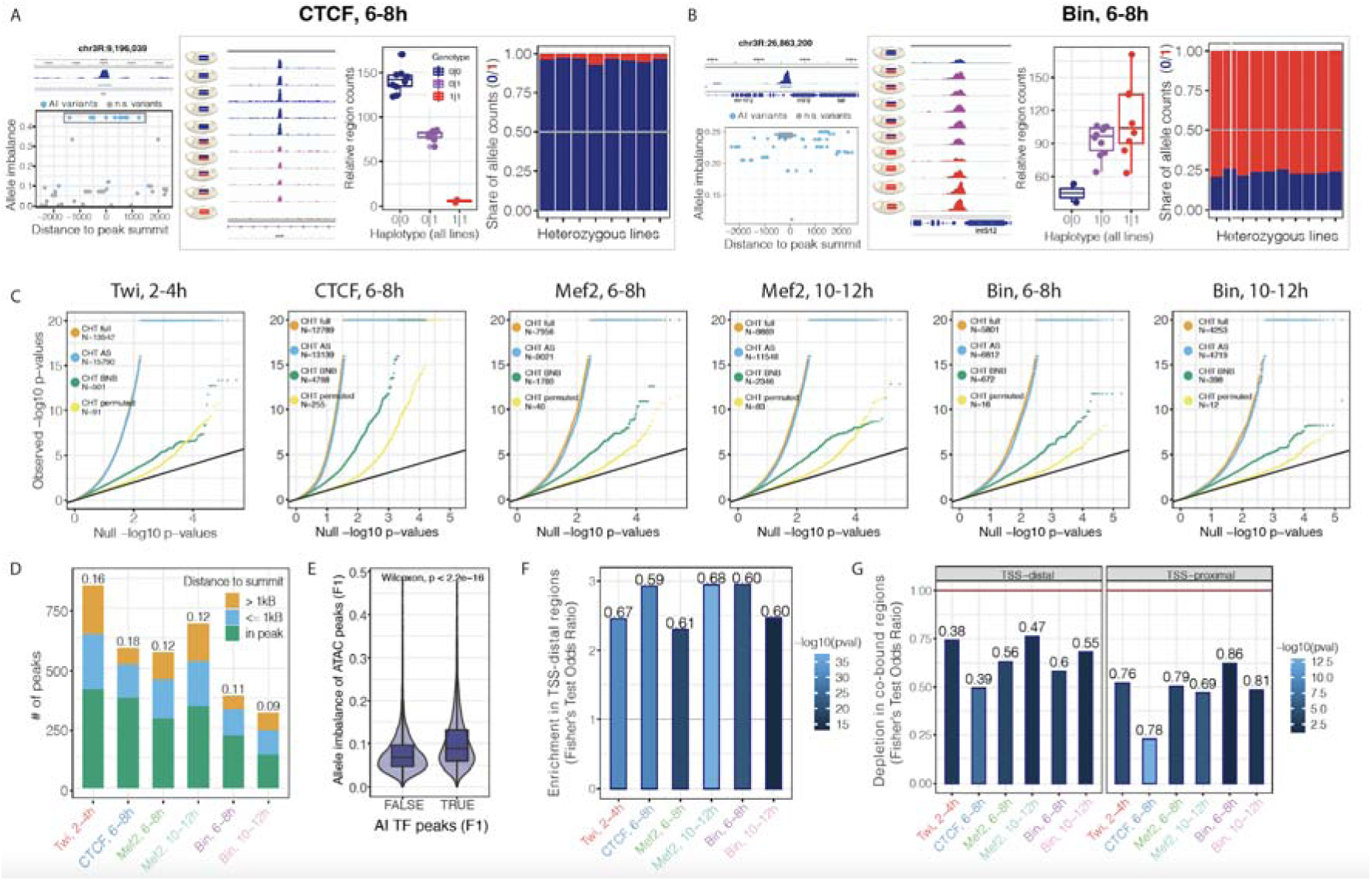
Genetic variation has extensive effects on TF binding during embryogenesis. **A.** Example of CTCF peak at 6-8h affected by genetic variation based on CHT. Panel on the top-left shows browser track for the corresponding peak (average signal across all lines). Coordinate of the peak summit is indicated on the top. Bottom-left panel shows allele imbalance (y-axis) for all variants in 2.5 kB radius around peak summit. Significant variants are shown in blue. For a subset of significant variants with the same genotype (grey box on bottom-left panel), browser tracks for all lines and two components of CHT are shown (three panels on the right). Schematic embryos next to browser tracks indicate genotypes of the lines. Components of the CHT are: normalized total read counts for the corresponding peak in all lines (total read depth, left) and share of reads mapped to the reference and alternative alleles in heterozygous lines (allele ratios, right). Colours represent genotypes: blue (reference), red (alternative), magenta (heterozygous). **B**. Same as A. for the imbalanced peak of Bin at 6-8h. **C**. QQ-plots showing CHT results for Twi at 2-4h, CTCF at 6-8h, Mef2 at 6-8h, Mef2 at 10-12h, Bin at 6-8h, and Bin at 10-12h. For each dataset, actual p-values are plotted against uniform p-value distribution for the full CHT (orange), allele-specific (AS) component of the CHT (blue), read depth (BNB) component, and CHT with permuted genotypes (yellow). Number of variants significant at 1% FDR with AI > 0.1 are provided in the legend. **D.** Number of AI peaks (y-axis) per condition (x-axis). AI peaks are defined as peaks with at least one associated significant variant. Shares of AI peaks in the total number of peaks with genetic variation are shown over the bars. Colors represent location of the top variant (significant variant with the lowest p-value per peak) relative to the peak summit. **E**. Average allele imbalance of ATAC-seq peaks overlapping AI and non-AI peaks from our dataset. Allele imbalances of ATAC-seq peaks were quantified for the same F1 crosses in (Floc’hlay *et al*, 2021). **F**. Enrichment of AI peaks in TSS-distal regions per condition (x-axis). Fisher’s test odds ratios (AI vs. non-AI peaks) are plotted on y-axis. Numbers over the bars indicate share of TSS-distal peaks in all AI peaks. Color represents p-value (-log10). **G**. Depletion of AI peaks in the regions co-bound by at least two TFs from our dataset for TSS-distal (left) and TSS-proximal (right) AI peaks. Fisher’s test odds ratios (AI vs. non-AI peaks) are plotted on y-axis. Numbers over the bars indicate share of co-bound AI peaks in all AI peaks. Color represents p-value (-log10). **E**. Consensus peaks overlap for each TF with the indicated number of genetic variants (0, 1-20, 21-50, 51-80, >80 variants) within 5 kB regions centred on peak summits.

Many of the identified significant variants were co-localized, and likely genetically linked. After grouping variants by TF peaks, we identified between 9% (Biniou 10-12h, 313 peaks) and 18% (CTCF 6-8h, 581 peaks) of TF peaks that are significantly affected by genetic variation (referred to as AI peaks, Fig. 2D). In 45-65% of AI peaks, the top significant variant (minimum p-value per peak) was inside the peak, while the other 35-55% of peaks were presumably affected by more distal genetic variants up to 2.5 kB away from the TF peak summit (Fig. 2D). Between 31-50% of AI peaks had INDELs among the significant variants, the majority of which were short (only 5-9% of AI peaks had INDELs longer than 10 bp). Allele imbalance associated with INDELs was slightly stronger compared to SNPs in most conditions (Supplementary Fig. 3B), although the difference was surprisingly low and only significant for Twist at 2-4h, Mef2 at 6-8h, and Biniou 10-12h (Kruskal-Wallis, p-value < 0.01). Our measurement of allelic imbalance in TF binding are further supported by AI in chromatin accessibility (Fig. 2E, Supplementary Fig. 3C), using ATAC-seq measurements in embryos from the same genetic crosses (Floc’hlay et al, 2021a). Furthermore, AI TF peaks are strongly enriched in putative enhancers (TSS-distal regions; Fisher’s test odds ratio 2.2-2.9, Fig. 2F), extending previous findings for open chromatin and H3K27ac (Floc’hlay et al. 2021). Both TSS-distal and proximal AI peaks are depleted in regions co-bound by the other TFs tested here (Fisher’s test odds ratios 0.24-0.72; Fig. 2G), as well as by other TFs (using a large collection of ChIP data from the ModERN database (Kudron et al. 2018) (Supplementary Fig. 3D) and from ubiquitously accessible DHS (Supplementary Fig. 3E). This suggests that the effects of genetic perturbations are buffered through redundant binding of other TFs to the same element e.g. through the collaborative action of TFs to maintain open chromatin. These findings agree with recent observations from single cell ATAC data in *Mef2* mutant embryos (Secchia et al. 2022) and suggest that the effects of genetic variation are more likely to be detected in genomic regions with less regulatory redundancy.

Overall, our results demonstrate that genetic variation has an extensive impact on TF occupancy *in vivo* during embryogenesis, even for essential developmental regulators. The effects of genetic variants on TF binding likely depend on the genomic context and complexity of the regulatory region, with promoter regions (TSS-proximal) and co-bound regions being less affected.

### Allele-specific binding preferences reconstruct functional TF motifs via *in vivo* ‘saturation mutagenesis’

To assess the extent to which AI in TF binding is due to genetic variants (SNPs or INDELs) within the cognate TFs’ binding site (TFBS), we scanned positional weighed matrixes (PWMs) of Twist, Mef2, Biniou and CTCF in 30 bp windows centred on all quantified variants (475,102 autosomal variants) in both the reference and alternative alleles (Methods). This identified a disruption of the cognate TFBS in 16-33% of AI peaks (Fig. 3A, i.e. AI peaks with motifs), which is significantly higher than expected by chance using a matched background of non-AI peaks (Supplementary Fig. 4A). Another 33-44% of AI peaks have a significant variant outside but in direct proximity to (within 100 bp) the TF’s motifs, which in similar to what is expected by chance (Fig. 3A, Supplementary Fig. 4A). When variants inside TF motifs were not associated with AI, the corresponding motifs were likely false positives of the motif prediction, as they were more frequently located either outside the peak (Fig. 3B) or further away from the peak summit (Fig. 3C) compared to motifs disrupted by significant AI variants. This indicates how AI in a tightly controlled system can pin-point the likely functional motif within or in the vicinity of a ChIP peak. Overall, all TFs had strong enrichment of AI variants inside their cognate motifs within their ChIP peaks (Fisher’s test odds ratios 2.1-3.2, Supplementary Fig. 4B, left), while no enrichment outside the ChIP peaks was observed, attesting to the specificity of the approach (Supplementary Fig. 4B, right). No motifs of other TFs from the CIS-BP database were significantly enriched in AI variants, when variants in the cognate TF motifs were excluded (Supplementary Table S3).

**Figure 3.**
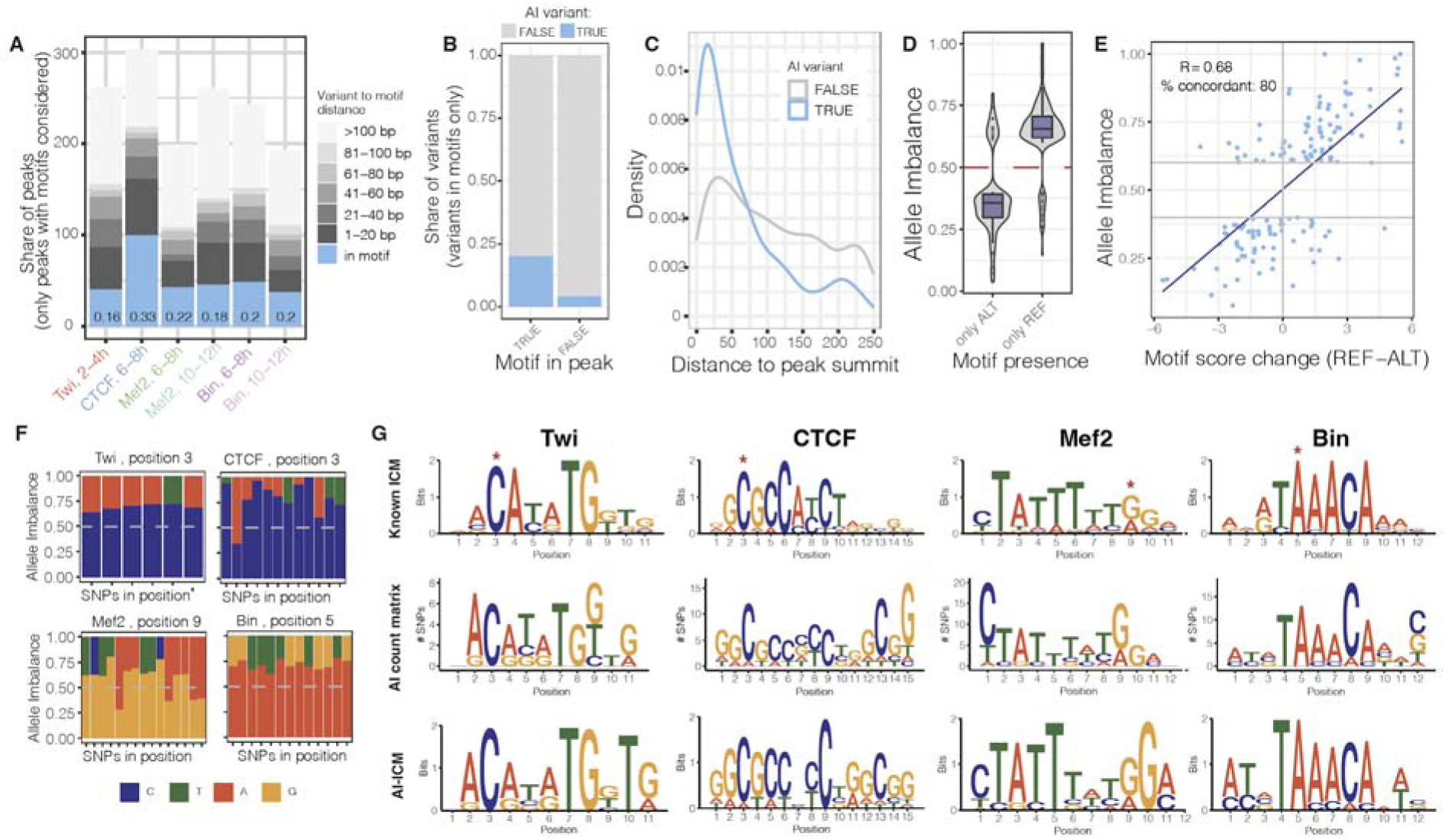
Allele-specific binding provides functionalised TF motif binding preferences at single base pair resolution: **A.** Distribution of AI peaks by the shortest distance of significant variants to TF motif (only AI peaks with motifs considered). Shares of AI peaks with significant variants disrupting TF motif are indicated for each condition (blue color and numbers). **B**. Share of significant (blue) and non-significant (grey) variants among all variants in motifs located in and outside TF peaks (x-axis). Combined data for all conditions is shown. **C**. For significant (blue) and non-significant (grey) variants in motifs, distribution of variants’ location relative to peak summit. Only motifs in peaks are considered. Combined data for all conditions is shown. **D**. Allele imbalance (y-axis) of peaks with significant variants in motifs, when motif is called only in the reference or only in the alternative allele (x-axis). Combined data for all conditions is shown. AI above 0.5 indicates reference allele bias. **E**. Peak AI (y-axis) vs motif score change (reference – alternative allele, x-axis) for significant variants inside TF motifs. Only cases when motif is called in both alleles are considered. Combined data for all conditions is shown, variants with absolute motif score change below 1 are not shown. Share of concordant variants (variants for which motif score change agrees with direction of allele imbalance) and Pearson correlation coefficient (R) between AI and score change are indicated. **F**. Allele imbalances associated with variants disrupting TF motifs: Twi at position 3, CTCF at position 3, Mef2 at position 9, and Bin at position 5. Each bar represents a single SNP, colors represent two alleles for each SNP, and y-axis shows allele imbalance. **G**. Motif logos of analyzed TFs generated from known PWMs (top row) or constructed based on AI SNPs in motifs (middle and bottom rows). Logos in the middle row represent counts of preferred nucleotides at each position, which were then transformed into information content matrixes and visualized as standard motif logos (bottom row). ICM – information content matrix.

As the AI is an experimental measure of the quantitative effect of genetic variants on TF binding, we next assessed if the measured differences in TF binding between alleles are in agreement with the differences in binding affinity as defined by PWM scores. This is akin to a saturation mutagenesis analysis, measuring the impact of all genetic variants within a TF’s motif on the TF’s binding, across multiple endogenous instances of the factor’s motif, in an *in vivo* chromatinized context. When the TFBS is completely ‘mutated’ to the extent that it is below the detection threshold in the alternative genotype, and thus only present in the reference genotype, the associated peak has very strong reference allele bias, and vice versa (Fig. 3D). Remarkably, when the motif is called in both genotypes, the associated allele imbalance is highly correlated with the changes in motif PWM scores (Pearson correlation coefficient R=0.68, Fig. 3E). Despite the relatively low number of variants overlapping TF binding motifs (i.e. not reaching saturation), most positions within motifs had SNPs associated with AI, thus allowing us to quantify the changes in occupancy (ChIP signal) when each motif base is altered. For example, we identified 6 SNPs disrupting position 3 of the Twist motif at different locations in the genome, and in all cases allele C was preferred for binding over the alternative alleles A and T (AI 0.6-0.7, Fig. 3F), in agreement with the Twist consensus motif (Fig. 3G, top row). Similarly, in all 13 AI SNPs at position 5 of the Biniou motif, allele A was preferred for occupancy over T and G (AI 0.65-0.75, Fig. 3F). We summarised these results by generating AI-based PWMs for all TFs based on the frequencies of their preferred alleles for occupancy at each position (Fig. 3G, Methods). Reassuringly, these ‘genetic’ based motif logos are very similar to the affinity-based PWMs for all TFs, which are often generated from *in vitro* binding assays on naked DNA (Fig. 3G, top vs. bottom rows). Thus, allele-specific binding information can confirm and extend the functional importance of each base within TF binding motifs at single-nucleotide resolution. This ‘functional’ assessment of the importance of each base will only improve in studies where more genotypes are assessed.

### Concordant changes in allelic imbalance between TFs and stages indicates a common regulatory mechanism

Although we found many variants linked to AI directly in the TF’s motif (discussed above), this can explain only a fraction (16-33%) of all AI peaks with a motif, as also observed in previous studies in yeast and human (Ding et al. 2014; Reddy et al. 2012; Kasowski et al. 2010). This implies widespread indirect effects of genetic variation on TF binding, including the disruption of motifs for a second factor that is cooperatively binding with the first TF. To identify such cases, we searched for AI peaks for two or more TFs that were co-affected by the same genetic variant, taking advantage of the extensive co-binding amongst the TFs profiled.

Cases where the same variant was significantly associated with pairs of AI peaks across time or between TFs were relatively rare: 16,0% (172/1,075) of Mef2 peaks and 12,2% (76/621) Biniou peaks at two time-points, and 5,8% (822/14,053) of pairs of different TFs (of 13 TF:time combinations) (Fig. 4A). This suggests that some variants impacting Mef2 and Biniou binding do so in a time-point specific manner, leading to significant allelic imbalance only at either 6-8h or 10-12h of embryogenesis. This is in keeping with our previous study on condition-dependent eQTLs during embryogenesis, where the main variants (many in putative enhancers) impacted gene expression only at one specific stage (Cannavò et al, 2016b). All co-affected peaks had high correlation in their AI (Fig. 4A-D, Supplementary Fig. 5A-F), which holds true for the same TFs at different time-points (0.93 for Mef2 and 0.97 for Biniou), for pairs of mesodermal TFs (0.76-0.95), and even between the insulator protein CTCF and mesodermal factors (0.75-0.95). Variants overlapping the co-affected peaks (green dots on Fig. 4B,C and Supplementary Fig. 5A-F) were almost exclusively concordant in the direction of their AI (both reference or both alternative bias), while there were only few variants that showed opposite AI effects on co-affected peaks, which were mostly coming from proximal and non-overlapping peaks (orange and blue dots). When restricting the analysis to the most significant variants for both peaks in the pair, the number of co-affected peaks was lower but the correlation increased for most pairs, especially for the same TF across two time-points: 0.97 for Mef2, and 0.99 for Biniou, across the two time-points (Supplementary Fig. 5G).

**Figure 4.**
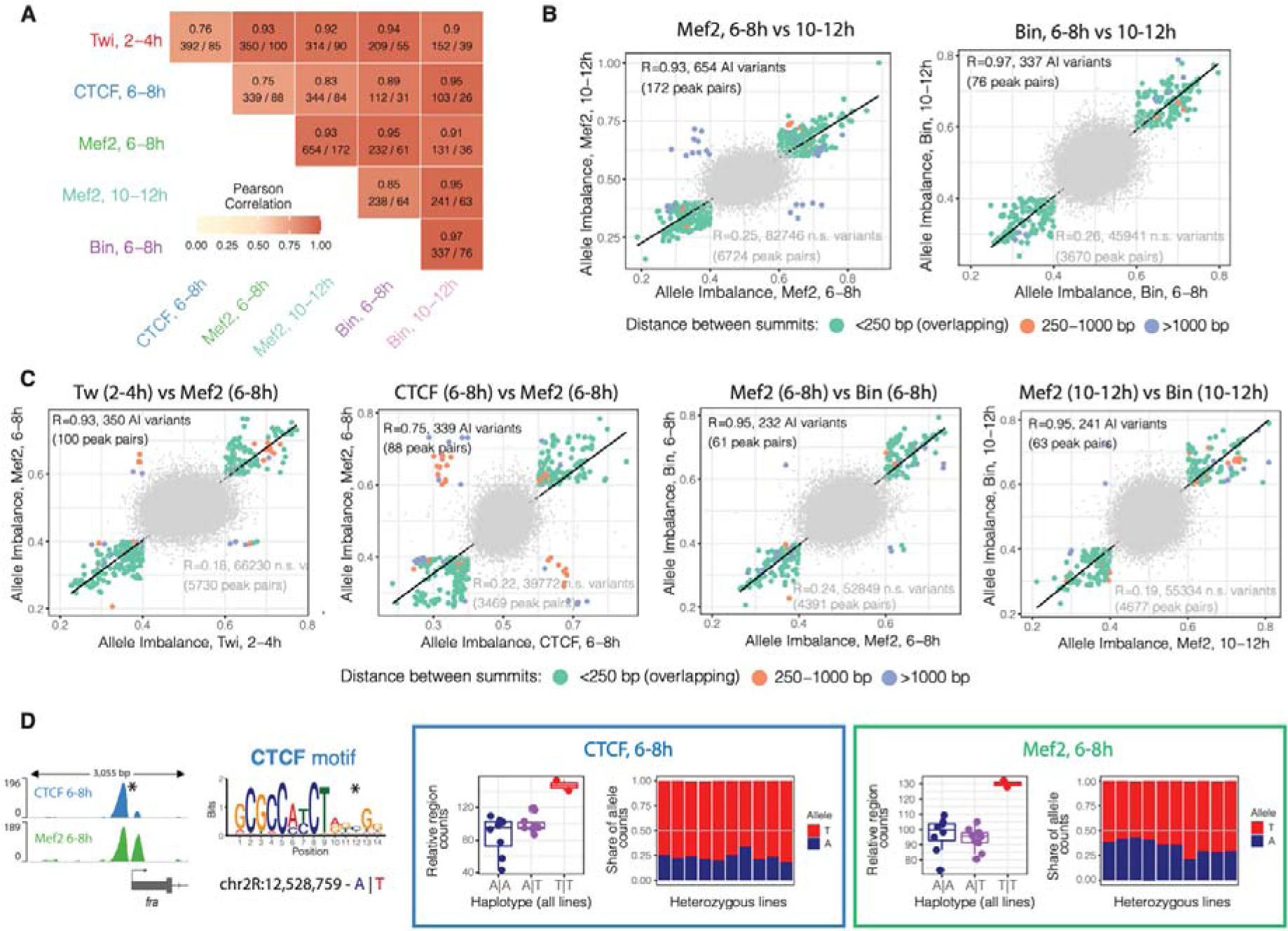
Variants outside cognate TF motifs might act through disrupted cooperative and collaborative binding: **A.** Correlation between AIs of variants in co-affected peaks for different conditions. Pearson correlation coefficient, number of significant shared variants / number of co-affected peaks are indicated. Only variants with unique AI values are considered for each pair of co-affected peaks. **B-C.** Correlation in AI for variants affecting pairs of conditions: Mef2 at 6-8 vs. 10-12h and Bin at 6-8 vs. 10-12h (B), Twi at 2-4h vs Mef2 at 6-8h, CTCF vs Mef2 at 6-8h, Mef2 vs Bin at 6-8h, and Mef2 vs Bin at 10-12h (C). Each coloured dot represents a significant variant (all significant variants with unique AI values per peak considered). Colour represents distance between co-affected peaks: overlapping peaks (green), less than 1kB between summits (orange), 1-2.5 kB between summits (blue). Grey dots represent all non-significant variants in both conditions. Spearman correlation coefficients are shown (top left). The number of variants and co-affected peaks are indicated for significant variants (top-left) and non-significant variants (bottom-right). **D**. Examples of a pair of co-affected peaks for CTCF and Mef2 at 6-8h where one of TF motifs was disrupted by variants. Left: browser tracks showing co-affected peaks (merged across all genotypes per TF) and affected TF motif (affected position marked with asterisk, reference and alternative alleles are indicated in blue and red, respectively). Boxes represent coverage counts for CTCF (blue) and Mef2 (green) split by allele. Left: normalized total read counts for the two genotypes for each of the two co-affected peaks. Right: allelic ratios (from CHT) in all heterozygous lines. Replicates are included as independent observations.

In one example, both CTCF and Mef2 binding at 6-8h are impacted at the promoter region of the *fra* gene (Fig. 4D, left), which share 47 significant variants, including 30 variants with minimum p-value for both TFs (adjusted p-value = 0), highlighting the challenge in identifying the causal variant. In this locus, one of those 30 variants (A/T, chr2R:12528759) disrupts a CTCF motif at position 12 resulting in the motif being called only in the alternative allele, even though the affected base in the motif has low information content (Fig. 4D, left). Both TFs have consistent alternative allele bias (Fig. 4D, right) suggesting that the CTCF motif disruption might lead to AI of a mesodermal TF (Mef2). However, overall, a surprisingly small fraction of co-affected peaks were associated with the disruption of either TFs’ cognate TFBS (2-12 peak pairs for different TF combinations, Supplementary Table S4), suggesting more complex cooperativity or collective binding, or perhaps effects on accessibility that then blocks the binding of any factor. Moreover, the majority of co-affected peak pairs had multiple associated variants (32 significant variants on average, 22 significant variants for peak pairs with TF motif disruptions), which further complicates the identification of causal variants even when they affect TF motifs.

Overall, our results indicate that the effects of genetic variants on co-affected TF peaks are highly concordant, suggesting the disruption of cooperative or collaborative binding. However, identifying the underlying mechanisms is hampered by the large number of highly significant linked variants and requires further prioritisation of significant variants.

### Basenji prioritises causal variants, contextualises variant effects and reveals mechanisms of co-recruitment

TF peaks with AI are linked to a median of 9 variants (range 1 - 121) due to residual linkage disequilibrium (LD). To disentangle LD and elucidate the potential functional variants, we trained a deep neural network model, Basenji (Kelley et al. 2018), to predict the effect of each variant separately. For training we used an extensive dataset of ChIP-seq bound regions during all stages of the *Drosophila* life cycle, and DNAse regions during embryogenesis, to learn bound regions within the wildtype reference *Drosophila* genome. The input training data included 1,205 ChIP-seq datasets from ReMap 2022 (Hammal et al, 2022a), which we reprocessed to produce RPGC normalised and background subtracted bigwig files, when a matching ChIP input was included in the database (Methods; Supplementary Table S6); 19 DNAse Hypersensitivity tracks from whole embryos and FACS purified muscle and neuronal cells (Reddington et al. 2020) and 6 ChIP-seq tracks from this study obtained by merging F1 crosses and replicates within a condition, giving a total of 1,230 ‘occupancy’ tracks. The reference genome was separated into 1,041 non-overlapping tiled sequences with a length of 131 kb. We randomly separated 214 sequences for the validation dataset, 193 sequences for the test dataset, and the remaining 634 (∼60%) sequences were used for training (Methods). Basenji performs very well in recovering the coverage of all tracks on the validation sequences (Fig. 5A,B), with a median Pearson correlation of 0.74 for ReMap 2022, 0.70 for DNAse and 0.78 for this study’s merged ChIP-seq data (Fig. 5B). Basenji can therefore learn the features of accessible or bound chromatin and is capable of reliably predicting the coverage of TF ChIP-seq tracks in *Drosophila*.

**Figure 5.**
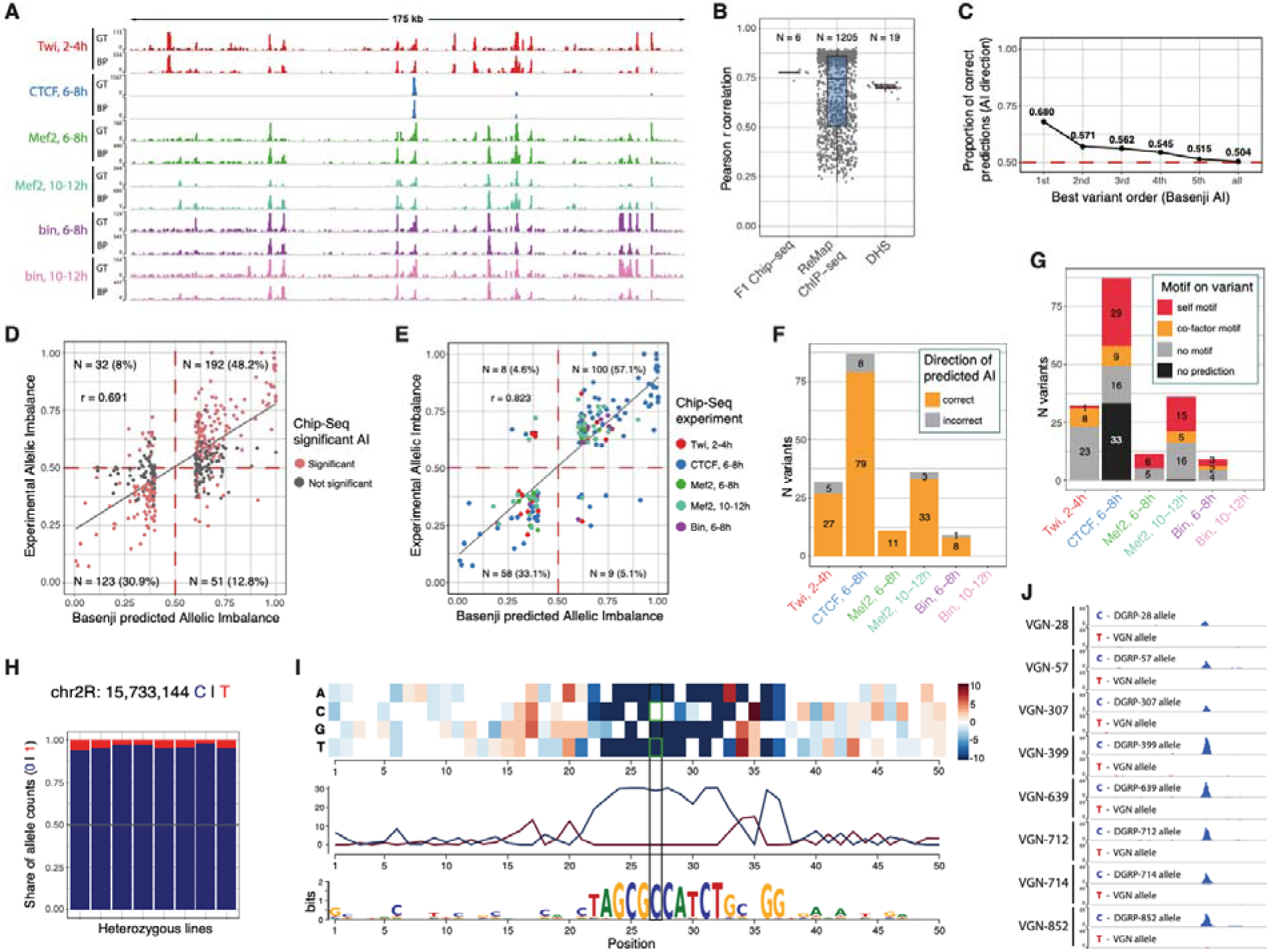
Basenji prioritises causal variants and predicts disrupted motifs via saturation scores: **A.** Ground truth (GT) coverage from merged F1 crosses compared with Basenji predicted (BP) coverage on test sequences from chromosome 2L. **B.** Pearson R correlation between ground truth and Basenji predicted tracks on test chromosome 2L for all targets used in training (ReMap n=1205, DHS n=19, F1 ChIP-seq n=6). **C.** Fraction of correct predictions (same direction of pAI and experimental AI) for variants within the same peak, ranked by absolute pAI from Basenji predictions. This test includes all peaks associated with at least one variant with significant experimental AI. **D.** Correlation between predicted AI and experimentally measured AI for variants with the highest absolute Basenji AI per peak (absolute pAI > 0.1). 79.1% of Basenji predictions are in the correct direction with a Pearson R = 0.691. **E.** Correlation between strong Basenji pAI (absolute pAI>0.1) and strong experimental AI (absolute AI>0.1). 90.2% of predictions are in the correct direction. Colours correspond to ChIP-seq samples. **F.** Counts of strong predictions per ChIP-seq sample divided by correct (same direction of effect; orange) and incorrect (grey) AI predictions. **G.** Same as in F) but samples are coloured by motif predictions overlapping the causal variants from saturation scores. Presence of cognate motif overlapping the causal variant (red), predicted cofactor motif overlapping the causal variant (orange), no motif overlapping the causal variant (grey) and failed saturation score prediction (black). **H.** Proportion of allele specific reads mapping to reference (C) or alternative (T) alleles on variant chr2R:15733144 for CTCF 6-8h. **I.** Saturation scores around variant chr2R:15733144 correlating with high AI in CTCF 6-8h sample, reveals that Basenji predicts a CTCF motif that is disrupted by a C to T variant. **J.** Visualization of ChIP-seq signal separated by maternal and paternal allele on the CTCF peak affected by variant in panel I. All DGRP lines harbour the C allele while the virginizer line has the T allele.

We then used the trained Basenji model to predict the effect of variants on TF binding. Variants were scored for their predicted effect on ChIP-seq signal, allowing to prioritise variants that caused AI. We generated reference and alternative sequences centred around each variant with experimental AI. Employing the trained model, we predicted the ChIP-seq peak signal for the reference and alternative sequences and computed a predicted AI (pAI) score at the variant position. pAI is defined as the predicted coverage on the reference allele divided by the sum of the predicted coverage on reference and alternative alleles. Thus, pAI is entirely based on the genomic sequence around the variant and independent of LD.

We assessed whether pAI prioritises variants within the same ChIP-seq peak. For this, we ranked significant variants (from the CHT full test) associated to the same ChIP-seq peak based on their absolute pAI value, and then checked the proportion of correct AI predictions in each rank (pAI and AI>0.5 or pAI and AI<0.5) (Fig. 5C). Looking at each peak’s top ranked variants, the pAI correctly predicts the direction of experimental AI in 68% of cases, indicating that the top pAI variants are enriched for causal variants. This proportion increases to 79.1% for top-predicted variants with large predicted effect size (absolute pAI>0.1, Fig. 5D) and decreases to background for lower ranked predicted variants. Overall, this suggests that pAI is a powerful tool to prioritise causal variants among those associated with the strongly imbalanced peaks.

Next, we explored how pAI can prioritise variants with unknown effects. The top pAI variants that had significant experimental AI, predicted the correct direction of effect in >90% of cases (Fig. 5E). The set of top pAI variants that were also the top experimental AI are enriched at (i) TSS-Distal peaks (130/175, Hypergeometric p=4.25e^-21^), (ii) being within cognate ChIP-seq peak (174/175, Hypergeometric p=1.72e^-34^), and (iii) overlapping the cognate TFBS (86/175, Hypergeometric p=3.83e^-90^) (Supplementary Fig. 7A) compared to all top pAI variants including those with no experimental AI. Interestingly, they are not evenly distributed across TFs, with CTCF representing half of these cases. The complexity of the CTCF motif (14 bp with high information content) and the strong relationship between its motif presence and binding (Ramírez et al. 2018), might increase the power of Basenji’s predictions for this factor. This suggests that the ability of sequence-based models to predict the impact of genetic variation in TF binding will vary depending on the type of DNA binding domain of the TF, which has important implications for future studies. To investigate the molecular mechanism of the 175 strong pAI predictions, we applied *in silico* saturation mutagenesis (Lee et al. 2015; Kelley et al. 2016) to uncover base-pair specific importance scores and reconstruct relevant motifs around these variants (Fig. 5E). For the majority (68% (119/175)), the model predicts at least one motif within 75bp of the strong pAI variants (Supplementary Fig. 7B) indicating that pAI scores are generally based on their effect on known motifs. Top pAI variants without known motifs are more likely incorrect predictions compared to top pAI variants without any predicted motif (Hypergeometric p=0.015) (Supplementary Fig. 7C), indicating that meaningful predictions of the regulatory landscape surrounding the variant is crucial for Basenji’s performance. A variant overlapping a cognate CTCF motif, for example, has a very strong effect on CTCF binding with an AI of 0.925 (Fig. 5H) and impacts the central part of the cognate motif leading to a strong pAI (0.997) (Fig. 5I). The reference C allele is present across all DGRP lines while the alternative T is only seen in the maternal line. By separating reads overlapping the variant we can recover the allele specific coverage that shows a striking AI at this locus (Fig. 5J).

Among the 175 variants, 7 have an effect on 2 different TFs or stages of embryogenesis and in all cases the direction of effect is the same between conditions, and pAI could correctly predict the AI. This suggests collaborative or cooperative binding. Different TFs were influenced by this to varying extents, again likely reflecting differences in the manner in which they are recruited to DNA. For CTCF and Mef2, the majority of top pAI variants overlaps their cognate motifs, rather than another TF’s putative co-operative motifs: CTCF pAI variants correspond to 29 cognate versus 9 potential co-factor motifs, Mef2 pAI to 21 cognate versus 5 co-factor motifs. This includes four variants associated with changes in Mef2 binding at both 6-8h and 10-12h of embryogenesis. This indicates that in the majority of cases, these two TFs do not bind cooperatively, and rather depend on the presence of their own motif. However, their occupancy (and motif presence) can influence the occupancy of other factors. In contrast, Twist appears highly cooperatively - only one significant motif variant (1/8) was in the cognate Twist motif, while the others were in potential cofactors motifs within the same ChIP-peaks. This includes significant variants in a Twist AI peak at 2-4h and also in either CTCF, Mef2 or Biniou AI peaks at 6-8h. In all cases the experimental AI for Twist and the putative co-factor’s binding have concordant directions, suggesting that they bind cooperatively (Fig. 5G, Supplementary Fig. 7F-H, and Supplementary Table S5). Although we lack F1 Chip-seq data for CTCF and Mef2 at the corresponding time-point as Twist (2-4h), we verified that both CTCF (Pollex et al. 2022) and Mef2 (Zinzen et al. 2009) co-bind with Twist at 2-4h at the respective regions, indicating that Twist binding is facilitated by CTCF and Mef2 acting as co-factors. Interestingly, in all cases the TFs’ binding at the later time-point has a stronger AI suggesting a downstream effect across embryogenesis (Supplementary Fig. 7F-H). An interesting example is a T to C variant with pAI impacting both Twist binding at 2-4h and Biniou binding at 6-8h, which does not impact either Twist or Biniou motifs, but is positioned in between the two motifs (Supplementary Fig. 7D-E). This may represent an example where flanking sequences can impact TF affinity, for example by altering DNA shape (Yella et al. 2018; Rohs et al. 2009), and (compatible with this) the sequence between the two motifs might be important for cooperative binding. To explore all saturation mutagenesis results we created a Shiny app accessible at http://furlonglab.embl.de/data.

### AI uncovers mechanisms of lineage specific recruitment of CTCF

Although the majority of AI impacting CTCF is due to variants within the CTCF motif (29 pAI from Basenji), our results uncovered a number of examples where CTCF binding is impacted by variants in a lineage specific TFs’ motif, including 9 predicted co-factor motifs. We uncovered, for example, three cases where variants in the Mef2 motif are associated with altered Mef2 and CTCF binding at 6-8h, and 5 cases where variants in the Twist motif impacts Twist binding at 2-4h, and CTCF binding at 6-8h (Supplementary Fig. 6B). For CTCF, 6-8h was the only stage that we performed the ChIP-seq at – however as the TF is expressed earlier, it is very likely that this SNP would also lead to AI in CTCF binding at 2-4h, or alternatively Twist binding early in embryogenesis may act to prime this site for CTCF binding at later embryonic stages. CTCF is ubiquitously expressed, while Twist and Mef2’s expression is tissue specific, restricted to the mesoderm and/or its differentiating muscle. This indicates that tissue specific TFs may play a role in recruiting CTCF (co-operatively) to a subset of sites. CTCF binds to the majority of its sites across all tissues (invariant peaks), matching its ubiquitous expression, in both *Drosophila* and mice (Pollex et al. 2024; Behera et al. 2018). However, there is a small fraction of tissue specific CTCF peaks, which our genetic data measuring AI indicates are recruited cooperatively by tissue specific TFs. Similar findings were also observed in mouse erythroid cells (Behera et al. 2018), suggesting that this is a common feature.

## DISCUSSION

In this study, we used F1 embryos from crosses of inbred *Drosophila melanogaster* lines to profile allele-specific binding of three essential developmental TFs (Twist, Mef2, Biniou) and one insulator protein (CTCF) at three embryonic time-points. This tightly controlled genetic design proved powerful in quantifying AI especially considering the small sample size (8 crosses). Our results demonstrate that variants with *cis* effects on TF binding (AI peaks) are common, affecting 9-18% of TF peaks in our dataset. This is in the range observed in previous studies in the context of human lymphoblastoid (Kilpinen et al. 2013; Tehranchi et al. 2016) and erythroid (Behera et al. 2018) cell lines or in a tissue (mouse liver) in a differentiated homeostatic state (Wong et al. 2017). The fact that we see a similar extent of variation for factors that are essential for embryogenesis, being influenced by genetic variants in the context of embryogenesis is surprising. Due to the large genetic diversity in our F1 design, most positions within each TF’s motif were covered, allowing us to recover the functional importance of each base pair within the motif. Variants in motifs are associated with strong AI, and allelic preferences at these positions are highly concordant with TF binding affinity scores based on PWMs.

Controlling for mapping biases is crucial for measuring AI in a reliable way. At the same time, INDELs prove difficult to assess due to mappability issues as they create sequence shifts between alleles. We expanded the WASP pipeline to include reads overlapping short INDELs both at the mappability filters and AI tests. Including INDELs improved our results by increasing mapped reads by 13-28%, leading to a substantial increase in the number of AI TF peaks that we could detect. As expected, functional variants are enriched for INDELs (5.8% more likely to be significant compared to SNPs) making them especially relevant to include. The functional impact of INDELs was surprisingly similar to SNPs – the change in TF binding was only slightly stronger (Supplementary Fig. 3B), suggesting that INDELs with strong phenotypic effects are relatively rare. This is counterintuitive, and suggests that INDELs with strong effect sizes are most likely selected against in these inbred *Drosophila* lines, since all lines are homozygous viable, while *twist*, *Mef2* and *biniou* are essential for embryonic development and viability.

### The ability to associate genetic variation to molecular phenotypes (AI) differs depending on the regulatory region and TF involved

Genes that are impacted by genetic variation are enriched or depleted for genes with distinct biological functions – TFs and developmental genes are generally depleted, while genes involved in metabolism are enriched (Floc’hlay et al. 2021; Cannavò et al. 2016). This is in part due to an ascertainment bias, as genetic variants impacting TF or other essential genes’ are often embryonic, or even cell, lethal, but it could also be due to more inherent buffering of regulatory elements for TFs (and other developmental regulators) compared to metabolic genes (Floc’hlay et al. 2021; Sigalova et al. 2020). Within the genes that have AI, different regulatory regions also have different sensitivities to the functional impact of genetic variation. TSS proximal regions (promoters) are less affected than TSS distal (putative enhancers) regions. As TSS-proximal regions are generally characterized by highly accessible chromatin, which is relatively invariant across tissues and developmental stages compared to distal elements (Reddington et al. 2020), the impact of genetic variants may be more buffered in promoter regions resulting in less changes in TF binding. This is in line with observations that promoters are genetically more conserved than distal regions (Cheng et al. 2014) and confirms our previous findings examining the impact of genetic variation on open chromatin and H3K27ac, which showed that allelic variation is both more frequent and has greater magnitude at distal regulatory elements (putative enhancers) compared to promoters, despite genetic variation itself being more common at promoters (Floc’hlay et al. 2021).

Even within regulatory elements, different TFs appear to be impacted differently by genetic variation, which likely reflects differences in the way they interact with DNA. For example, Basenji could learn the rules for sequence changes impacting CTCF binding much better than the other three TFs tested. All four TFs have different types of DNA binding domains, and therefore different bases are directly contacted by the TF. While the other three factors have motifs of typical length (6-8bp), CTCF’s motif is much longer and of high information context, and CTCF has a higher retention time on DNA (in the minute range) compared to most other TFs. It may also reflect how cooperative a TF is – we found many examples of variants that impact the binding of multiple TFs. Interestingly, this included variants impacting the motif of tissue specific TFs (Mef2) that also impacted the binding of CTCF, which is a ubiquitously expressed TF. The extent of genetic variation impacting two or more TFs is underestimated here due to the sample size, and is likely much more extensive than currently envisaged. Such differences in the ability to model the impact of genetic variation for different TFs has important implications for sequence-based models going forward, and will remain a future challenge.

### The mechanism of some genetic variation impacting TF binding remains inaccessible

While Basenji performed very well, the strongest predictions are biased towards variants in motifs within the peaks with the strongest AI. This could be because most causal variants are actually overlapping motifs (cognate or co-factor), with the model being inherently capable of highlighting them, since the first layer of filters can be traced to motifs. Being completely unbiased towards existing motif knowledge, the model can also recognize unknown motifs and interpret surrounding sequence context, such as DNA shape, and could uncover potential cases of cooperative binding. However, Basenji could only make meaningful predictions for a fraction (5,2%, 175/3,368) of significant AI peaks. Purely correlative methods (such as the Combined Haplotype Test applied here) are much more sensitive in identifying variants associated with AI peaks, but would likely over-represent variants outside peaks and motifs that are linked to causal variants in LD. On the other hand, a subset of the genetic variants outside the TF’s peak are likely causal and not linked to LD. We were not able to resolve the mechanism of that variation here (Basenji’s predictability appears to decrease at increasing distances from a TF motif/peak), and perhaps it requires more sophisticated models or the integration of additional types of data such as 3D chromatin proximity such as Hi-C or Capture-C data.

In conclusion, pAI offers an orthogonal approach to prioritize causal variants and interpret their effect on TF binding. However, to understand the full spectrum of genetic variation other methods are required to uncover the hidden mechanisms of more distal variation on TF binding and gene expression.

## METHODS

### Overview of the dataset

Crosses were generated between a single isogenic maternal line (VGN) and eight paternal DGRP lines (DGRP-28, DGRP-307, DGRP-399, DGRP-57, DGRP-639, DGRP-712, DGRP-714, DGRP-852). ChIP-seq was performed on embryos from 8 F1 and 2 parental lines (VGN and DGRP-399), each in two biological replicates, for the following TFs and embryonic time-points: Twist at 2-4h, CTCF at 6-8h, Mef2 at 6-8h and 10-12h, Biniou at 6-8h and 10-12h (6 conditions × 10 lines x 2 replicates = 120 samples in total).

#### *Drosophila* genetics and embryo collection

To obtain F1 hybrid embryos, males from eight genetically distinct inbred lines from the DGRP collection (MacKay et al. 2012) were crossed to females from a “virginizer” (VGN) line. The VGN line contains a heat-shock-inducible pro-apoptotic gene (*hid*) on the Y Chromosome (Starz-Gaiano et al. 2001) of a laboratory reference strain (*w1118*) that kills all male embryos after a 37°C heat-shock.

Freshly hatched adults were placed in embryo collection containers with standard apple cap plates. For ChIP-seq experiments, following three 1 h pre-lays, the flies were allowed to lay embryos for 2h. The embryos were then aged for 2h, 6h or 10h to reach the developmental intervals 2-4h, 6-8h or 10-12h. The embryos were collected and dechorionated using 50% bleach, and washed with deionized water (dH2O) and PBT (phosphate buffered saline (PBS) containing 0.1% Triton X-100). Dechorionated embryos were crosslinked in 1.8% formaldehyde for 15 min at room temperature. Fixation was stopped by the addition of PBT + Glycine 125mM and embryos were washed with PBT. After drying the embryos on tissue, they were snap-frozen in liquid nitrogen and store at −80°C until use.

### Chromatin preparation and ChIP-seq on *Drosophila* embryos

Nuclei were extracted from the embryos using a 7ml tissue homogenizer (Wheaton) and HB buffer (15 mM Tris pH 7.4, 0.34 M Sucrose, 15 mM NaCl, 60 mM KCl, 0.2 mM EDTA, 0.2 mM EGTA). After douncing 20 times with the loose and 20 times with the tight pestle, nuclei were spun down at 2.465 xg for 7 min at 4°C and then washed one time with HB buffer and one time with PBT (1x PBS, 0.1% Triton X-100). Nuclei were then counted using the BD LSRFortessa™ Cell Analyzer, centrifuged at 2.000g for 10 min at 4°C and the supernatant was removed. Nuclei were snap-frozen and stored at −80°C until use.

The frozen nuclei pellet was resuspended with an appropriate amount of RIPA buffer (140mM NaCl, 10 mM Tris-HCl pH 8.0, 1 mM EDTA, 1% Triton X-100, 0.1% SDS, 0.1% Na-deoxycholate, 1x Roche Complete Protease inhibitors; < 40 million nuclei: 1 ml; 40-100 million nuclei: 1.5 ml; >100 million nuclei: 2 ml) and incubated on ice for 10 min. The nuclei were then sonicated using a Bioruptor Pico in 15ml sonication tubes with sonication beads (Diagenode C01020031) according to the manufacturer’s instructions for 15 cycles (30 sec on / 30 sec off). The supernatant was transferred into 1.5 ml low binding tubes (Eppendorf, 0030108051) and centrifuged at 20.000 xg for 10 min at 4°C. Aliquots of chromatin were snap-frozen in liquid N2 and stored at −80°C.

For quality control of the chromatin, one aliquot of 30 μl was RNase treated (50 mg/ml final) at 37°C for 30 min and reverse cross-linked using a final concentration of 0.5% SDS and 0.5 mg/ml Proteinase K overnight at 37°C for 10 hours, and 65°C for 8 hours on a thermomixer with interval shaking. The next day the DNA was purified with Phenol-Chloroform purification and precipitated with ethanol, Sodium Acetate pH 5.3 and glycogen to obtain pure DNA. The size distribution was assessed by gel electrophoresis by running 100 ng of DNA on a 1.2% agarose TAE gel. The majority of the DNA was concentrated between 250 and 500 bp. The concentration was measured using Qubit hs DNA (Thermo Fisher Scientific, Q33230).

ChIP-seq was performed as described in (Bonn et al. 2012), using antibodies directed against Twist (Sandmann et al. 2007), Mef2 (Sandmann et al. 2006) and Biniou (Jakobsen et al. 2007) generated in our lab, and against CTCF from R. Reinkawitz’s lab. Antibodies were incubated overnight with chromatin in RIPA buffer in a total volume of 900 μl using 3 μl of rabbit anti-Mef2, 1 μl guinea pig anti-Twist, 1 μl rabbit anti-Biniou, and 1 μl of rabbit anti-CTCF. The next day 25 μl of magnetic protein A/G beads (Dynabeads, Invitrogen, 10002D and 10004D) were washed with 1ml of RIPA buffer and added to the IPs for an additional 3 hour incubation on the rotating wheel at 4°C. The ChIPs were then washed for 10 min on the rotating wheel with 1x 1 ml RIPA, 4x 1 ml RIPA-500 (500 mM NaCl, 10 mM Tris-HCl pH 8.0, 1 mM EDTA, 1% Triton X-100, 0.1% SDS, 0.1% Na-deoxycholate), 1x 1 ml LiCL buffer (250 mM LiCl, 10 mM Tris-HCl pH 8.0, 1 mM EDTA, 0.5% IGEPAL CA-630 CA-630, 0.5 % Na-deoxycholate) and 2x 1 ml TE buffer (10 mM Tris pH 8.0, 1 mM EDTA) in the cold room. The chromatin was then RNase-treated and reverse cross-linked as described for the quality check of the chromatin.

A molecular barcoded ChIP-seq library was prepared with all ChIPed-DNA obtained using the NEXTflex^™^ qRNA-seq^™^ Kit v2 (HiSS Diagnostics, 520999-03). The quality of the libraries was assessed on a Bioanalyzer (Agilent), and libraries displayed a peak around 350-600 bp. For each ChIP, at least two completely independent biological replicates were performed. ChIP-seq libraries were paired-end sequenced with 75bp for ChIP samples and 150bp for ChIP inputs using an Illumina NextSeq 500 platform at the EMBL Genomics Core Facility.

### *De novo* genotyping of parental lines

*Drosophila* fly lines have a short reproduction cycle leading to hundreds of generations within a few years. This causes accumulation of *de novo* variants creating divergence from original genotyping data, bottleneck effects, and further homozygosity – all of which can also occur in established cell lines. To avoid the need for downstream filtering (gDNA correction) to achieve balanced coverage between parental chromosomes, as used in previous studies (Floc’hlay et al. 2021), we performed deep *de novo* sequencing of the parental lines and created *de novo* genotyping of the exact lines used for the experiment to avoid mappability biases from unidentified variants.

We collected 10 adult flies from each of the parental lines used in this study were collected and snap frozen: DGRP-28, DGRP-307, DGRP-399, DGRP-57, DGRP-639, DGRP-712, DGRP-714, DGRP-852 and VGN. To extract the genomic DNA, the flies were crushed with a handheld pestle until powdered, while kept frozen into liquid nitrogen. 500 μl buffer A (Tris-HCl pH 7.5 100 mM, EDTA pH 8.0 100mM, NaCl 100mM, SDS 1%) were added and mixed with the powdered flies. 20 μl of PK proteinase K (PK) 20 mg/ml were added and the samples incubated for 3h at 65°C. PK was inactivated by incubation at 95°C for 10 min. The samples were then treated with 1 μl RNAseA 10 mg/ml and incubated for 30 min at 37°C. DNA was extracted with 500 μl phenol/chlorophorm/IAA, followed by 500 μl chlorophorm. The aqueous phase was transferred to a new tube and 3 volumes of 100% ethanol were added. The pellet was washed with 1 ml 70% EtOH and air-dried and resuspended in 25 μl 10mM Tris-HCl. 2 μg of gDNA per sample were diluted in 100 μl of TE buffer. The gDNA was sonicated with a Bioruptor Pico using 6 cycles of 30s ON and 90s OFF. 1 μg of sheared gDNA was used for library preparation with the NEBNext Ultra II DNA library kit. We enriched for fragments of 300-400bp of length and amplified the library with 3 PCR cycles. The sequencing was performed using an Illumina NextSeq to obtain 150bp paired-end reads.

The lines were sequenced deeply giving between 54M and 92M reads to achieve a whole genome coverage that ranged between 74X and 125X. Variant discovery was performed following the GATK Best Practice Workflow for short variant discovery. Reads were trimmed using TrimGalore (TrimGalore) (v. 0.5.0) with options ‘-q 30 --phred33 --fastqc --illumina --length 75 –paired’. Reads were then mapped to a joint genome of *Drosophila melanogaster* (dm6) and Wolbachia (AE017196) to quantify the parasite contamination using bwa mem (Li & Durbin, 2009) (v. 0.7.17) with options ‘-T 20’. Reads were then sorted using samtools (Li *et al*, 2009) (v. 1.9) and duplicates were first marked with picard (Broad Institute 2015) MarkDuplicates (v. 2.16.0), then removed together with reads not mapped in proper pairs using samtools with option ‘-F 1548 -f3’. We then applied GATK (McKenna et al. 2010) (v. 4.1.0.0) base recalibration on the mapped reads using as a reference the variants included in the full DGRP catalogue from 205 inbred fly lines as a reference (Huang et al. 2014) with GATK BaseRecalibrator and GATK ApplyBQSR. To increase variant discovery and improve line to line comparisons we performed a joint variant call using GATK HaplotypeCaller ‘--min-base-quality-score 20 -G StandardAnnotation’. Variants were then filtered using two sets of cutoffs to produce a hard stringent filtered and a more lenient filtered variant setss list corresponding to a high confidence and a high sensitivity variant lists, respectively. The hard stringent filtered variant set was obtained using bcftools (Li et al. 2009) (v. 1.9) with options ‘MQ > 58 & MQRankSum > −2,5 & MQRankSum < 2,5 & QD > 20 & SOR < 1,5 & FS < 10 & ReadPosRankSum > −4 & ReadPosRankSum < 4’ while the lenient filtered variant set was obtained using options ‘MQ > 40 & MQRankSum > −12,5 & MQRankSum < 12,5 & QD > 2 & SOR < 3 & FS < 60 & ReadPosRankSum > −8 & ReadPosRankSum < 8’.

### ChIP-seq read mapping and mappability filter

FASTQ files with ChIP-seq reads were pre-processed by removing unique molecular identifiers (UMIs) using JE v.1.2 (Girardot et al. 2016) with parameters *LEN=8 BPOS=BOTH ADD=TRUE SAME_HEADERS=TRUE XT=1 ZT=0 RCHAR=’:*’ and removing adaptor readthrough with skewer v0.2.2 (Jiang *et al*, 2014) with parameters *-m pe -z -x ‘NNNNNNNNNAGATCGGAAGAGCACACGTCTGAACTCCAGTCACNNNNNNNNATCTCG TATGCCGTCTTCTGCTTG’ -y ‘NNNNNNNNNAGATCGGAAGAGCGTCGTGTAGGGAAAGAGTGTAGATCTCGGTGGTCGCCGTATCATT’*.

Low quality bases were removed with seqtk trimfq v1.2 (https://github.com/lh3/seqtk) with default parameters. Reads were mapped to the reference dm6 genome with bwa v0.7.15 (Li and Durbin 2010) using bwa mem with option ‘-M’. BAM files were sorted and indexed with picard SortSam (Broad Institute, 2015). Reads were filtered requiring a minimum mapping quality of 10. One input sample (VGN-DGRP307 at 2-4h) was removed from the analysis due to failed QC. For the peak calling, input samples were merged across all lines per time-point (see below).

Since the ChIP-seq samples came from genetically diverse lines of *Drosophila melanogaster,* mapping reads to the reference genome could result in some reads being mapped better to the reference allele (reference bias). To correct for such potential mapping biases, we adapted the original WASP code from (Van De Geijn et al. 2015) to include INDELs into the mappability correction (the original version discarded reads with indels). WASP-INDEL is available on http://furlonglab.embl.de/resources/tools.

In brief, all mapped reads that overlapped at least a SNPs or INDEL in the parental *Drosophila* lines (from the *de novo* genotyping, lenient variant set) were identified. Alleles in those reads were substituted with all possible allele combinations from the parental VCF file (up to 64 different allelic combinations per read, default) and mapped back to the reference genome with the same options as specified above. If all combinations of the reads overlapping variants mapped with a mapping quality > 10 uniquely to the same position of the reference genome as the original read, they were retained for subsequent analysis. Finally, duplicate reads were removed with Je suite (Girardot et al, 2016) using random filtering of duplicates to avoid reference bias as discussed in (Van De Geijn et al. 2015).

### ChIP Peak calling and Irreproducibility Discovery Rate (IDR) analysis

We implemented an IDR pipeline (Li et al. 2011) as recommended in ENCODE guidelines (Landt et al. 2012). ChIP peaks were called using MACS2 (Zhang et al. 2008) with relaxed p-value threshold (0.5) for each replicate/pseudoreplicate ‘-p 0.5 --format BAMPE --gsize 142573017’. Input files used in peak calling were merged per time-point (across all lines) using samtools merge v.1.9 (Li et al. 2009). IDR tool v2.0.4.2 was used (https://github.com/nboley/idr). The number of peaks at IDR 1%, 2% and 5% (global IDR value) were recorded (Supplementary Table S2) from the following steps:

– Between true biological replicates (Nt)
– Between pooled pseudoreplicates (Np)
– Between self-pseudoreplicates of biological replicate 1 (N1)
– Between self-pseudoreplicates of biological replicate 2 (N2).

The majority of samples showed high reproducibility and satisfied the following quality criteria suggested for the IDR analysis: (1) Nt and Np within a factor of 2 and (2) N1 and N2 within a factor of 2 (Supplementary Table S2). At IDR 1%, two samples for one line (VGN-DGRP714 for Mef2 and Bin at 10-12h) showed a very low number of reproducible peaks between replicates (Nt=130 and 51, respectively) due to a failed replicate 1 (N1=354 and 117, respectively), and were therefore excluded from all downstream analysis. Based on this QC analysis, we included reproducible peaks at IDR 1%, and required a peak to be present in at least three fly lines to be included in the consensus peak sets (see below).

### Defining consensus peak sets

Consensus peak sets were constructed using DiffBind package in R (Ross-Innes et al. 2012). For each condition (TF + time-point), we used reproducible peaks between true replicates (‘conservative’ peak set) at 1% IDR in each line. The consensus set was produced with dba.count function using TMM normalization (score=DBA_SCORE_TMM_READS_FULL), fragment size from bam files (fragmentSize = 0) and control reads were normalized by relative library size (default). Only peaks present in at least three lines were considered (minOverlap = 6 since we have 2 replicates for each line). For the resulting consensus set, summits were recalculated based on read coverage, and consensus peaks were resized to 500 bp around the new summits (summits=250).

### Phased genotypes for F1s

Phased genotypes for the stringent F1 variant set were constructed with a custom script (construct_phased_vcf_with_replicates_with_gl_tabix.py) using genotypes of parental lines. Only variants without missing information that were homozygous in all nine parental lines were considered (1,799,462 variants considered, 96% of all 1,878,415 variants). In addition, we kept only bi-allelic variants for subsequent analysis (1,735,077 variants, 96% of the above).

### Total and allele-specific counts per target regions

Allelic counts for each variant in the final VCF file were calculated using get_counts.py script from WASP-INDEL. Variants in 2.5 kB radius of consensus peak summits (target regions) were defined as test variants (get_target_regions.py script with parameters: min_read_count=400, min_as_count=50, min_het_count=1, min_minor_allele_count=1). For each test variant and target region, total and allele-specific read counts were calculated with get_region_data.py script. Script update_total_depth.py from the original WASP pipeline (Van De Geijn et al. 2015) and used to adjust total counts for the GC content and the fraction of reads in peaks.

### Combined haplotype test

The combined haplotype test (CHT) from the WASP suite (Van De Geijn et al. 2015) was used to define variants associated with differences in read depth or/and allelic imbalance among individuals. CHT was run separately for each of the 6 biological conditions (Twist at 2-4h; Mef2, CTCF and Biniou at 6-8h; Mef2 and Biniou at 10-12h). To increase the power of the test, we treated biological replicates as individuals and included both parents (only containing homozygous positions) and F1s (both homo- and heterozygous positions). At 10-12h, line VGN-DGRP714 was removed (see above) resulting in the set of 18 samples (20 samples at 2-4h and 6-8h). Only variants on autosomes were considered for the CHT. We ran CHT, allele-specific (Beta Binomial) and read depth (Beta Negative Binomial) parts of the CHT on real data and permuted genotypes. P-values from all these tests were plotted against expected p-value distributions (Fig. 2C).

### Quantifying variant effects

Adjusted p-values (FDR, BH correction) were calculated on the sets of variants quantified by CHT, an FDR threshold of 0.01 was used to define significant variants. Reference allele bias was defined for each variant as 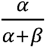, where *α* and *β* are expected counts for reference and alternative alleles (from CHT), respectively. Allelic imbalance (AI) was defined for each variant as 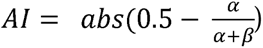, where *α* and *β* are defined above, and 0.5 is the expected allelic ratio in case of no imbalance. A threshold of AI=0.1 was used to define strongly imbalanced significant variants. Relative region read counts were defined as 100 * (total region read counts) / (expected region read counts), where expected region read counts were estimated with update_total_depth.py script from the WASP suite.

### Motif analysis

*De novo* motif discovery and central enrichment was performed on the top 1000 peaks from the consensus peak sets from each condition (ranked by average TMM scores across all samples) using MEME-ChIP v 4.11.3 (Bailey et al. 2009) with second order of the Markov background model (-order 2). This was complemented by scanning for known TF motifs in the consensus peak sets (separately for each condition) and 30 bp intervals centred on all quantified variants (both reference and alternative allele) using fimo 4.11.3 (Bailey et al. 2009) on both strands (default) with nucleotide frequencies as background (calculated with fasta-get-markov v. 4.11.3, from MEME suite). We used optimized PWMs for Twist, Mef2, and Biniou (Zinzen et al. 2009), while all other PWMs were downloaded from CIS-BP database (Weirauch et al. 2015) version from 05.03.2019.

For all quantified variants, the distance to the nearest motif was recorded. To quantify the distribution of AI peaks by variant to motif distance (Fig. 3A), we selected the closest significant variant to the cognate TFBS for each peak. To construct a background set for each condition (Supplementary Fig. 4A), we used the following procedure: (1) from non-AI peaks with motifs, randomly select the same number of peaks as in AI peak set, (2) from the selected peak set select the number of associated variants equal to the number of significant variants, (3) for each peak, record minimum distance of variants to motif. This procedure was repeated 100 times, and the average number of peaks was reported for each distance interval.

For reconstructing motif logos from allele-specific binding (Fig. 3F-G), we used all significant SNPs with AI>0.1 overlapping motifs of corresponding TFs. For each position in the motif, we selected nucleotides that were preferably bound by the TF based on allele-specific information, e.g. selecting reference allele if *α*>β for the considered SNP and alternative allele otherwise. For each TF, we constructed positional frequency matrixes (PFM) based on the numbers of nucleotides preferentially bound at each position affected by variants. Resulting PFMs were then used as input to make PWM and seqLogo functions from the seqLogo package (Bembom and Ivanek 2024) in R with default parameters.

### ReMap transcription factor coverage tracks

The mapped bam files were processed according to the ReMap analysis pipeline (Hammal et al. 2022) and provided by Fayrouz Hamal and Benoît Ballester. According to the database annotation we were able to match 570 out of 1205 ChIP-seq samples with at least one corresponding ChIP input (same publication, conditions and sample of origin). In cases where multiple ChIP inputs were associated to the same sample, the bam files were merged with samtools merge (Li et al. 2009). All coverage tracks (ChIP samples and ChIP inputs) were generated using bamCoverage (Ramírez et al. 2016) with the following options “--binSize 10 --normalizeUsing RPGC --effectiveGenomeSize 125464728 --ignoreForNormalization chrX chrM chrY” with option “--extendReads 0” for single-end reads (no reads extention) and “--extendReads” for paired-end reads (reads were extended to match the fragment size). Samples with a corresponding ChIP input were input subtracted with bigwigCompare (Ramírez et al. 2016) with options “--binSize 10 --operation subtract”. All bigwigs are available for browsing and download at: http://furlonglab.embl.de/data.

### Basenji model and training

The input of the model is non-overlapping sequences from the reference dm6 genome. We extracted 1,041 non-overlapping sequences with a length of 131 kb across the genome, of which 214 sequences were randomly separated for the validation set, 193 sequences for the test set, and the remaining 634 sequences were used as a train set. With the fixed input size of 131,072 bp, for better learning 8,192 bp were cut out from both ends of the input and the training loss was calculated only on the centre crop. The centre crop then has a length of 114,688 bp. The output of the model comprises 1,230 tracks, of which 1,205 are derived from the ReMap tracks, 19 are DHS tracks, and the remaining 6 are from the F1 Chip-Seq tracks generated in this study.

During the training phase, we used random shift and reverse complement as data augmentation techniques. For the random shift, the fixed input size of 131,072 bp was reduced to 114,688 bp, and the centre of this crop was subjected to the random shift. The reverse complement randomly selected half of the training sequences in the mini-batch and replaced them with their reverse complement sequence.

The first part of the model is a convolutional block followed by a max-pooling operation with a pool size of 2. The convolutional blocks consist of the GELU activation function, convolution with a kernel of width 15 and 288 filters, and batch normalization. These convolution and pooling operations aggregate the base pair information into 128-bp bins. The convolutional block is repeated six times with a kernel of width 5 and increasing the number of filters from an initial 288 by 1.1776x each block to 768 filters by the end. Subsequently, 11 layers of residual blocks with dilated convolution are applied, followed by a final convolutional layer to make the prediction. The residual block comprises a GELU activation, a dilated convolutional layer with a kernel of width 3 and 384 filters, batch normalization, another GELU activation, a convolutional layer with a kernel of width 1 and the original number of filters, batch normalization, and dropout with a 0.3 rate. Each residual block features a skipped connection to the inputs of the block. The final output convolutional layer undergoes average pooling and a dense layer of size 1230 with softplus activation produces the model predictions.

The training of the model was performed with Stochastic Gradient Descent (SGD) with a learning rate of 0.1. The batch size was set to 4, and the momentum was configured at 0.99. Additionally, a clip norm of 2 was employed to control the gradient norm, preventing it from growing too large during training. The model training was executed on NVIDIA A100 Graphics Processing Unit (GPU). The code for training and evaluation of the model are available here: https://github.com/ZauggGroup/basenji_drosophila.

#### pAI and variants prioritization

For each variant, we created two sequences, reference and alternative allele, centred at the position of the variant. The reference allele contains the information from the dm6 genome and an alternative sequence introduces the variant in the sequence. The trained Basenji model was used to create predictions for the reference allele and the alternative allele individually. The pAI corresponds to the Basenji prediction for the reference allele divided by the sum of the Basenji prediction for the reference and the alternative alleles. If multiple variants had a significant association to the same peak (AI > 0.1 and FDR < 0.01), they were prioritized by ranking the pAI (|pAI - 0.5|).

#### Saturation mutagenesis

We performed saturation mutagenesis applying ‘basenji_sat_bed.py’ script within Basenji’s repository (Kelley et al. 2018). The saturation mutagenesis was calculated for the 6 conditions merged tracks (merge of crosses and replicates) that provide a baseline coverage. We used ‘sum’ as the prediction statistics and performed saturation mutagenesis 75bp upstream and downstream of the 176 variants that have both AI and pAI > 0.1. The saturation scores were then normalized on the average of all scores absolute values and the deltas were computed by subtracting the scores for the reference allele at each bp. The importance scores represent the sum of the absolute values of deltas per bp. We created the motif logos from the importance scores using R package motifStack (Ou et al. 2018).

## DATA AVAILABILITY

All raw data, which consists of xx demultiplexed files, were submitted to EMBL-EBI ArrayExpress (https://www.ebi.ac.uk/arrayexpress/browse.html) under accession numbers: E-MTAB-xxxx (F1 ChIP-seq) and MTAB-xxxx (DNA sequencing data for variant calling). The processed F1 bigwigs, reprocessed ReMap bigwigs, and variant calls are available on the Furlong lab web page, http://furlonglab.embl.de/data.

## ACKNOWLEDGMENTS

We thank all members of the Furlong and Zaugg labs for discussions and useful comments during the course of this study and on the manuscript. This work was technically supported by the EMBL Genomics Core facility and IT services, and financially supported by PhD Fellowship to F.H. from the Provence-Alpes-Côte d’Azur Regional Council (Région SUD), and ERC advanced grant agreement 787611 (DeCRyPT) to E.E.M.F.

## AUTHOR CONTRIBUTIONS

O.S, J.Z. and E.E.M.F. designed the study. R.R.V. and B.Z. performed the embryo collections and R.R.V. the ChIP-seq experiments. R.R.V. and M.F. performed the whole genome sequencing and M.F. performed the *de novo* genotyping. O.S. processed the F1 ChIP data and analyzed the results. A.R. and O.S. modified WASP for INDELs. F.H and B.B processed and provided the ReMap 2022 *Drosophila* ChIP-seq raw data, M.F. made input subtracted bigwigs. F.S. trained and evaluated the Basenji model, and M.F. analysed the model’s predictions with input from O.S. O.S., M.F. and E.E.M.F wrote the manuscript with input from all authors. E.E.M.F. funded the study. All authors discussed the results and commented on the manuscript.

